# Acidic Conditions Promote Clustering of Cancer Cell Derived Extracellular Vesicles and Enhance their Fusion with Synthetic Liposomes

**DOI:** 10.1101/2024.11.04.621963

**Authors:** William S. Fisher, Jessica Douglas, Sherwin Roshan, Ramon Perez, Sophia Wei, Logan Roberts, Kai K. Ewert, Cyrus R. Safinya

## Abstract

Extracellular vesicles (EVs) are endogenous vesicles secreted by cells. Exosomes, a subset of EVs ranging from 30 to 150 nanometers in diameter, contain cytosolic proteins, gene-silencing miRNA, and gene-encoding mRNA and have roles in intercellular communication. Exosomes show promise as cancer chemotherapeutic drug delivery vehicles given their low immunogenicity and cell-specific delivery of luminal contents to the cytosol of target cells. However, loading exosomes with cancer chemotherapeutic drugs is inefficient, which limits their therapeutic application. To overcome this barrier, methods have been developed which allow fusion of EVs with synthetic liposomes preloaded with therapeutic drug. While these methods show more efficient fusion than passive incubation of EVs with liposomes, they risk either damage to the membrane proteins of the EVs or contamination of the final EV-liposome hybrid with residual depletant molecules, which can cause side effects or hinder content delivery. Here, we present a new, weakly perturbative method which uses acidic conditions (pH 5) to significantly enhance the fusion of EVs and synthetic, neutral liposomes (NLs) compared to passive incubation in pH 7.4 at 37 °C. An adapted Forster resonance energy transfer (FRET) based lipid mixing assay confirms that fusion is enhanced with this method. This significant finding implies that lipid-only synthetic liposomes are able to fuse with EVs, creating EV-liposome hybrids under relevant temperature and pH conditions, with no other non-lipidic component, such as fusogenic amphipathic peptides, added to the synthetic liposomes. Remarkably, differential interference contrast (DIC) and fluorescence microscopy show that this enhancement of fusion corresponds with the onset of clustering of mixtures of EVs and NLs, or EVs alone, in acidic *but not neutral* pH conditions. The findings support a hypothesis that content release from EVs in early to late endocytic environment may be a combination of protein-protein clustering interactions and a lipidic component. Further, this study provides a novel method for enhanced fusion of EVs and liposomes which is expected to preserve EV membrane proteins and functionality towards the development of therapeutic hybrid drug delivery vehicles in nanomedicine applications.

## INTRODUCTION

Extracellular vesicles (EVs) are endogenous vesicles secreted by most cells with roles in intercellular communication. EVs are comprised of functionally distinct exosomes and microvesicles, which are differentiated by their size ranges, protein and nucleic acid content, and cellular pathways, which give rise to their formation [1, 2]. Exosomes (ranging from 30 nm to 150 nm in diameter) are capable of miRNA and mRNA transfer from donor to acceptor cells and alteration of gene expression and cell behavior in target (acceptor) cells, often in a cell-specific manner [3–7]. Exosomes have generated significant interest in the drug delivery field owing to their unique cell targeting and delivery capabilities.

The formation of exosomes begins with endocytosis, where the plasma membrane invaginates to form an intracellular vesicle (supplemental Figure S1). These new intracellular vesicles are transported to the early endosome (EE), which sorts extracellular material for final transport to specific cellular locations. A fraction of sorted material returns to the plasma membrane while the remainder is retained in the endosome as it matures into a late endosome (LE). The EE to LE maturation process corresponds to the acidification of its lumen by membrane-bound proton pumps and simultaneous invagination and pinching off of its membrane to form intraluminal vesicles (ILVs) containing cellular miRNA, mRNA and proteins. The majority of ILVs enclosed within the LE (referred to as multivesicular bodies, MVBs) are deposited into the highly acidic lysosome lumen (pH 4.8) and are enzymatically degraded following fusion of the MVB with the lysosome. However, a fraction of MVBs shuttle to and fuse with the plasma membrane to release their ILVs into extracellular space. These externalized ILVs are exosomes, which have inter-cellular communication functions that are not fully elucidated [2].

Exosomes circulate systemically, evidenced by their abundance in blood and urine [8], and have distinct, organ and cell specific accumulation based on their protein, lipid, and glycan enrichment patterns [1, 9, 10] imparting intrinsic targeting abilities [11]. Upon arrival at the target cell, exosomes are endocytosed and fuse with the membrane of cytosolic vesicles or the EE to release their contents into the cytosol [4, 12]. This fusion and content release mechanism is not understood but may use surface protein interactions which are promoted in the acidic conditions present in mature endocytic vesicles [4, 7, 13]. Select membrane lipids may also drive the transition of exosome membranes to a more fusogenic state [14].

The ability of exosomes to target specific cells in select cases and escape endosomal entrapment described above, combined with their observed capacity for hydrophilic and hydrophobic drug loading [12, 15–17], makes exosomes promising drug delivery vehicles complementing synthetic vesicles. Synthetic vesicles, or liposomes, are among the most investigated delivery vehicles for nucleic acid (DNA, siRNA, mRNA) and cancer drugs in therapeutic applications [18–27]. Numerous liposome formulations containing chemotherapeutic drug in their aqueous interior, including Doxil and Myocet, are in clinical use [28]. However, liposomal vectors face two major barriers to further clinical success: cell-specific targeting *in vivo* remains difficult to achieve, and entrapment of drug-loaded vectors in the endosome prevents complete therapeutic payload delivery [29, 30]. The huge success, with respect to delivery of the ionizable liposome-based mRNA vaccines against SARS-CoV-2 in the worldwide Covid-19 pandemic, is in part, due to the fact that they are injected directly into muscle; thereby, avoiding many of the challenges related to systemic targeted delivery *in vivo* that remain relevant, for example, in cancer therapeutics [31, 32].

For cationic liposomes carrying the chemotherapeutic drug paclitaxel (PTX), drug loading is a key factor in cytotoxic efficacy [22]. PTX is a hydrophobic chemotherapeutic with high potency that is used in the treatment of ovarian, breast, and non-small cell lung cancers [33–38]. The poor aqueous solubility of PTX requires that a carrier is used to deliver it to target cells in the body. Taxol®, a formulation of PTX in a 1:1 mixture of polyoxyethylated castor oil and ethanol, was the most prominent clinical carrier of PTX until 2010 [39]. However, Taxol induces severe PTX-independent hypersensitivity, which has led to extensive research to develop safer PTX carriers [34, 40, 41]. PTX encapsulated in the hydrophobic lipid bilayers of liposomes has emerged as a leading strategy for PTX delivery, with liposome encapsulated PTX formulations in late-stage clinical trials in the U.S. and Taiwan [42, 43]. Thus, exosomes are also expected to encapsulate PTX within their membrane interiors and a recent study reported that EVs were able to solubilize 15 to 20 μM PTX in aqueous media, greater than the 10 μM solubility limit of water, through passive loading [44]. To achieve meaningful PTX loading in therapeutics, new methods for the loading of PTX into exosomes must be developed.

An emergent strategy for drug loading and surface modification of EVs is the fusion of EVs with synthetic liposomes loaded with therapeutic drug [45–47]. With this strategy, the synthetic liposome can be functionalized with customizable targeting moieties on its membrane and loaded with therapeutic drug either in its lipid membrane, as with PTX, or aqueous interior. The customized synthetic liposomes are then fused with EVs to create hybrid vesicles which contain components of the liposomes and EVs and thus retain the biocompatibility of EVs while enhancing or modifying their cell specificity and drug loading capacity while transferring a therapeutic drug payload. Current methods to achieve this fusion include freeze-thaw cycling [46], passive incubation [48], polyethylene glycol (PEG) incubation [45], and extrusion of mixtures of liposomes and EVs [49]. Freeze thaw cycling, PEG-mediated fusion, and extrusion all facilitate more rapid fusion of EVs and liposomes than passive incubation. However, the use of freeze-thaw cycling and extrusion may cause damage of native EV membrane proteins and freeze-thaw cycling in particular can degrade encapsulated drugs [47]. PEG-mediated fusion, while effective in inducing fusion, results in PEG contamination of the final hybrid samples which can interfere with biological activity or drive PEG-mediated immune reactions [50]. Thus, further research is needed to develop liposome-EV fusion techniques which strike a balance between preventing damage to EV proteins and contents and effectively driving fusion.

The present study was designed to adapt a Forster Resonance Energy Transfer (FRET) based technique for tracking fusion of EVs and synthetic liposomes and interrogate a novel, *weakly perturbative* pH-based method for enhancing the fusion of liposomes and EVs with FRET and complementary techniques. To test existing and novel fusion methods for EV-liposome hybrids, EVs were isolated according to established protocols [44] and mixed with neutral liposomes (NLs). NLs consisting of zwitterionic 1,2-dioleoyl-*sn*-glycero-3-phosphoethanolamine (DOPE) and 1,2-dioleoyl-sn-glycero-3-phosphatidylcholine (DOPC) were developed which contained the labeled lipids 1,2-dioleoyl-sn-glycero-3-phosphoethanolamine-N-7-nitro-2-1,3-benzoxadiazol-4-yl (NBD-DOPE) and 1,2-dioleoyl-sn-glycero-3-phosphoethanolamine-N-lissamine rhodamine B sulfonyl (Rhodamine-DOPE). DOPE has an inverse cone shape and is known to promote membrane fusion [51, 52]. The labeled lipids are an established FRET pair used in lipid mixing assays which monitor the fusion of distinct lipid vesicles [45, 53]. In this system, the increase in NBD fluorescence is proportionate to the number and extent of vesicle fusion events. Vesicle fusion allows the diffusion of NBD and Rhodamine labeled lipid into empty membrane. Diffusion increases the average distance between the fluorophores to greater than 10 nm, diminishing the absorbance of photons emitted by excited NBD by Rhodamine.

Here, employing FRET we show that *lipid-only* synthetic NL liposomes are able to fuse with EVs, at 37°C and pH ranging from 7.4 to 5, with no other non-lipidic component, such as fusogenic amphipathic peptides, added to the synthetic liposomes. The FRET studies were conducted in reaction mixtures of NL liposomes and EVs isolated from human M21 melanoma and PC3 prostate cancer cells. A statistically significant enhancement of fusion was observed for mixtures of NLs and EVs in acidic conditions (pH 5) compared to neutral buffers at pH 7.4. Previous studies had only shown fusion between EVs and extracted plasma membranes; that is, between two biological membranes, each of which contained a heterogeneous mixture of lipids and membrane-proteins where fusion is expected to be driven, in part, by key proteins in each membrane [4, 13].

Remarkably, this enhancement of fusion at low pH and 37°C occurs concurrently with the onset of clusters of labeled EVs and mixtures of labeled EVs and NLs in acidic but not neutral conditions observed in optical imaging experiments employing differential-interference-contrast (DIC) and fluorescence microscopy. The possibility of a protein-membrane interaction, like the mechanism by which influenza virus hemagglutinin drives aggregation (via protein tethers) and promotes fusion of biological vesicles in acidic conditions [54, 55], is raised with these results. Interestingly, enhanced fusion of NLs and anionic liposomes containing DOPC and 1,2-dioleoyl-sn-glycero-3-phosphatidylserine (DOPS, −1e), to mimic the lipids with PS headgroups in EVs, was also observed in acidic conditions. Thus, the observed enhanced fusion of synthetic NLs and EVs in acidic conditions may contain, in part, a lipid-dependent mechanism.

The present study is expected to have significant impact on the development of EV-based nanotherapeutic drug delivery by enabling a gentle method for EV-liposome fusion and development of EV-liposome hybrids. The findings presented here suggest that acidic conditions promote the fusion of EVs and synthetic liposomes, without the need for accelerants which risk damaging native EV membrane proteins or contaminating the final sample. Thus EV-liposome hybrids can be created which retain the EV-derived machinery required for endosomal escape and cell-specific targeting while containing custom drug-loaded membrane and luminal drug contents introduced via synthetic liposome. Importantly, our FRET data shows that NBD-lipid and rhodamine-lipid molecules have been delivered from the NLs to the NL-EV hybrid vesicle created in the fusion process. Beyond their application to the formation of EV-liposome hybrid drug delivery vehicles, the finding that acidic conditions promote aggregation of EVs alone provides mechanistic insight on the formation and content delivery process used by EVs. An understanding of that process would have wide-ranging impact on nanomedicine by enabling endosomal escape strategies employed by native communication vesicles.

## MATERIALS AND METHODS

### Materials

Stocks of DOPE, DOPC, and DOPS were prepared from purchased powder (Avanti) dissolved in chloroform to 10 mM concentrations. Rhodamine-DOPE and NBD-DOPE pre-dissolved in chloroform to 0.768 mM and 1.082 mM, respectively, were purchased from Avanti. Polyethylene oxide (molecular weight average of 20 kDa) purchased from Sigma Aldrich was dissolved in water to 50 % (w/v). Clear polystyrene 96 well plates (Corning) were used for all lipid mixing assays.

### Liposome preparation

Solutions containing lipids and labeled lipids were mixed in chloroform:methanol (3:1, v/v) at 2mM (NLs) and 5 mM (anionic liposomes) for FRET assays and 1 mM for microscopy experiments. When included, NBD-DOPE and Rhodamine-DOPE were kept at 0.5 mol % of formulation composition. The chloroform:methanol solvent was evaporated under nitrogen gas stream for 7 minutes, then the lipid film was dried further under vacuum for 16 hours. The lipid film resulting was resuspended in high resistivity water (18 MΩcm) to the concentrations listed previously. Rehydrated films were then sonicated continuously for 7 minutes by a tip sonicator (Sonics and Materials Inc. Vibra Cell) set at 30 Watt output to form sonicated liposomes.

### Cell culture

The human prostate cancer cell line PC3 (ATCC number: CRL-1435) and melanoma cell line M21 (subclone of UCLA-SO-M21 derived from Reisfeld lab, Scripps Institute, La Jolla) were gifts from the Ruoslahti Lab (Burnham Institute, La Jolla). Cells were cultured in Dulbecco’s modified Eagle’s medium (DMEM; Invitrogen) supplemented with 10% v/v fetal bovine serum (FBS; Corning) and 1 % v/v penicillin and streptomycin (Gibco). Cells were grown in T75 flaks (Corning) at 37°C in a humidified incubator with 5% CO_2_ and split at a 1:5 ratio after reaching >80% confluence, at approximately every 48 hours, during maintenance and expansion.

### Extracellular vesicle isolation

Extracellular vesicles were isolated as described previously [44]. Briefly, FBS was depleted of native EVs by ultracentrifugation at 100,000 x*g* for 18 hours at 4°C and added to DMEM to make media with 10% EV depleted FBS. Then, 200mL of this EV depleted media was added to 20 flasks containing M21 or PC3 cells at 80% confluence and incubated for 48 hours to make conditioned media (CM). 200mL of this CM was taken for EV isolation. To clear cell debris and non-EV vesicles, e.g. apoptotic bodies, CM was centrifuged at 300 x*g* for 10 minutes, 2000 x*g* for 20 minutes, and 10,000 x*g* for 30 minutes, discarding the pellets each time. To isolate EVs, the resulting supernatant was centrifuged at 100,000 x*g* for 70 minutes and the pellets were resuspended in PBS, combined, and brought 20 mL PBS total in a single ultracentrifuge tube. After this washing step, the resuspended EVs in PBS were centrifuged again at 100,000 x*g* and the resulting pellet was resuspended in 500 μL of PBS.

### Nanoparticle tracking analysis (NTA)

Samples of EVs for NTA were made by diluting the unfiltered 500 μL EV pellet resuspension 1:1000 in PBS. Cationic liposomes (CLs) were prepared by mixing DOPC and 1,2-dioleoyl-3-trimethylammonium-propane (DOTAP) at a 1:1 molar ratio in chloroform:methanol (3:1, v/v) to 1 mM total concentration. The chloroform:methanol solvent was evaporated for 7 min under a nitrogen stream and further dried in a vacuum for 16 hours. The resultant film was resuspended in high resistivity water (18.2 MΩ cm) to the same 1 mM total concentration. This suspension was sonicated for 7 min with a tip sonicator (Sonics and Materials Inc. Vibra Cell, 30 W output) to form small unilamellar vesicles. This sample was diluted 1:1000 in deionized water. Nanoparticle tracking analysis on the diluted samples was performed using the NanoSight NS300 system (Malvern) and the associated NTA 3.0 analytical software (Malvern). Scattering mode was used for all acquisitions and acquisition and analysis settings were constant across measurements. These measurements were captured over 60 second intervals and repeated five times for each reported size versus concentration profile. There were 2858 ± 158, 4052 ± 178, and 1001 ± 104 valid tracks recorded for PC3 EVs, M21 EVs, and CLs, respectively. The measured concentration of particles at these dilutions were 8.88 × 10^8^ ± 0.44 × 10^8^, 10.21 × 10^8^ ± 0.34 × 10^8^, and 2.25 × 10^8^ ± 0.242 × 10^8^ for PC3 EVs, M21 EVs, and CLs, respectively. Measured concentrations of particles with diameters contained in each 5 nm bin from 10 to 1000 nm were adjusted by the dilution factor used in preparation of the NTA samples. Plots of diameter versus dilution-adjusted concentration for each sample were made in Python.

### Lipid Mixing Assay

Neutral liposomes (NLs) and anionic liposomes at 2 mM and 5 mM, respectively, and the desired lipid molar ratios were made as described above. NLs and either anionic liposomes or EVs were mixed in 150 μL high resistivity water, PBS, or sodium acetate buffer (100mM sodium acetate, 0.9 % (w/v) NaCl, pH adjusted by addition of HCl). NLs and anionic liposomes and NLs and EVs were mixed at the mol:mol or mol:particle ratios specified in the results and discussion section. These mixtures were prepared in a 96 well plate immediately before measurement using a Tecan M200 fluorescent plate reader. Fluorescence measurements used 460 ± 9 nm and 538 ± 20 nm wavelengths for excitation and emission, respectively, with a gain of 100, 4 readings per well, and a fixed plate z-position of 2100 μm. Temperature was held at 37°C for the extent of all lipid mixing assays and wells were mixed by automated linear and circular shaking before each measurement. When polyethylene oxide (PEO) was used, it was added after a single measurement of the vesicle mixture alone was taken to ensure that the rapid fusion at early timepoints did not mask the starting fluorescence. The average total fluorescence of each sample and fluorescence ratio for mixtures of EVs and NLs to unmixed NLs was calculated on Excel. The average and standard deviation of the fluorescence ratio from three repeat measurements in each condition were taken to determine statistical significance between conditions using the Student’s t-test in Excel.

### Microscopy

Neutral liposomes (NLs) and anionic liposomes at 1 mM and the desired lipid molar ratios were made as described above. NLs and either anionic liposomes or NLs and EVs were mixed at ratios of 1 nmol to 1 nmol or 1 nmol to 2×10^9^ particles in a final volume of 10 mL of high resistivity water, PBS, or sodium acetate buffer (100mM sodium acetate, 0.9 % (w/v) NaCl, pH adjusted by addition of HCl). EVs alone were imaged by diluting EVs 10-fold into either PBS or sodium acetate buffer adjusted to the listed pH. When used, PEO was added after combining NLs and EVs or anionic liposomes in solution and mixing. Immediately after mixing, 2 μL of each mixture was placed on glass slide within a parafilm cutout and covered by glass coverslip. Images were taken at either 20X or 40X magnification on an inverted Ti2-E microscope (Nikon) under DIC or fluorescence mode.

## RESULTS AND DISCUSSION

### Lipid mixing assay shows fluorophore dequenching corresponding with conditions which enhance vesicle fusion and correlates with PEO-induced clustering under fluorescence microscopy

To interrogate the extent of fusion between vesicles in aqueous solution, we employed a Förster resonance energy transfer (FRET) based lipid mixing assay with NBD and Rhodamine as the donor and acceptor fluorophores, respectively. In this assay, NBD and Rhodamine conjugated lipids were incorporated in the membrane of neutral liposomes (NLs) consisting of DOPC and DOPE known to promote membrane fusion [51, 52]. Fusion of a membrane containing NBD and Rhodamine conjugated lipids with a membrane lacking fluorophores leads to an increased average distance between NBD and Rhodamine fluorophores after lipid diffusion and results in increased total NBD fluorescence due to a weakened FRET effect [45, 53]. Therefore, the extent of fusion between NLs and vesicles with fluorophore deficient membranes is directly proportional to an increase in the ratio of NBD fluorescence observed in a mixture of NLs and fluorophore deficient vesicles compared to a sample containing only NLs. We express the data as a ratio of NBD fluorescence in a mixture of NLs and fluorophore deficient vesicles to NBD fluorescence in a sample of NL alone to account for changes in NBD fluorescence unrelated to membrane fusion such as photobleaching.

Under this theory of lipid mixing assay mechanics, an increase in the surface area of fluorophore deficient membrane available for NBD and Rhodamine conjugated lipid to diffuse into would increase the average distance between fluorophores and therefore increase the total NBD fluorescence relative to unmixed NLs, or the NBD ratio. The area of membrane available for fluorophore conjugated lipid to diffuse into increases with greater numbers of fusion events between NL and fluorophore deficient vesicles. Increased numbers of fusion events can be promoted by increasing the fusogenicity of the NL in a mixture of NL and fluorophore deficient vesicles at fixed amounts or by increasing the amount of fluorophore deficient vesicles relative to NL in the mixture. Figure 1 shows results which demonstrate that our assay responds in a manner consistent with this established theory. We mixed NLs with DOPE membrane contents ranging from 0 to 75 mol % and fluorophore deficient neutral liposomes composed solely of DOPC, termed DOPC-vesicles, in water at fixed lipid molar ratios R_DOPC-vesicle/NL_ (mol_DOPC (DOPC-vesicle)_/(mol_DOPC (NL)_ + mol_DOPE (NL)_)) = 9. The NBD ratio of the mixtures increased monotonically with the NL DOPE content (Figure 1A). This is consistent with the expectation that increased fusogenic lipid content should enhance the probability and extent of fusion on stochastic NL and DOPC-vesicle collisions.

**Figure 1.**
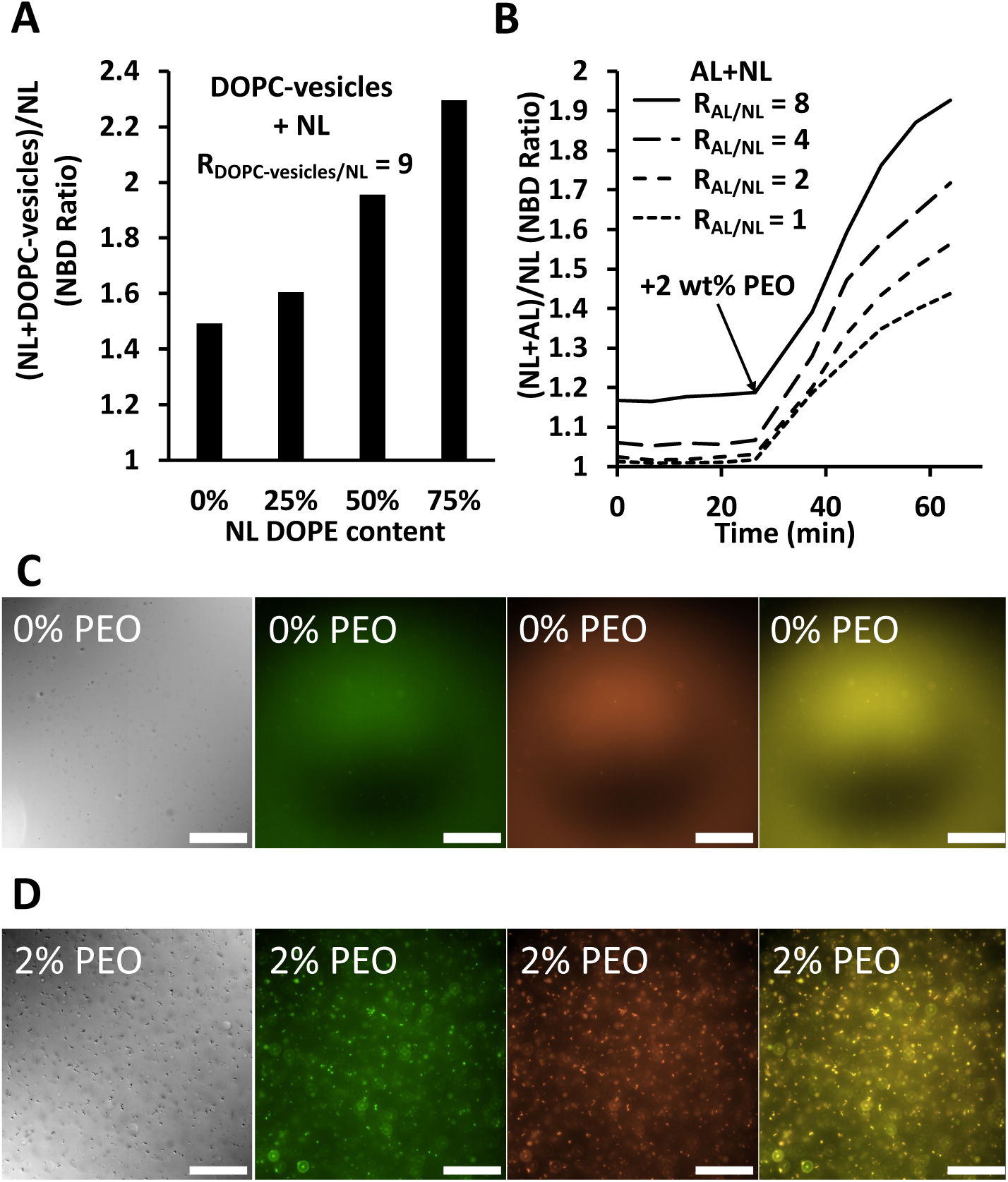
Total dequenched fluorescence (by FRET), differential-interference-contrast (DIC) and fluorescence microscopy, of mixtures of fluorescent neutral liposomes (NLs) and anionic or neutral liposomes. **(A)** Total normalized NBD fluorescence of NBD and Rhodamine containing NLs (DOPE_X_:DOPC_99-X_:NBD-DOPE_0.5_:Rho-DOPE_0.5_; X from 0 to 75 mol% DOPE) mixed with fluorophore deficient neutral DOPC vesicles in water at lipid molar ratio R_DOPC-vesicle/NL_ (mol_DOPC (DOPC-vesicle)_/(mol_DOPC (NL)_ + mol_DOPE (NL)_)) = 9. The total dequenched NBD fluorescence of these mixtures was taken after incubation for 2 hours at 37°C and normalized to the NBD fluorescence of a sample of only NLs. **(B)** Total NBD fluorescence of NBD and Rhodamine containing NLs at 50 mol% DOPE mixed with fluorophore deficient anionic liposomes (ALs, DOPC_90_:DOPS_10_) in water with R_AL/NL_ ((mol_DOPC (AL)_ + mol_DOPS (AL)_)/(mol_DOPC (NL)_ + mol_DOPE (NL)_)) increasing from 1 to 8 at 37°C. 20 kDa PEO was added after 26 minutes to a final concentration of 2 % (w/v), with total dequenching NBD fluorescence taken at multiple timepoints throughout and normalized to a sample of only NLs treated identically. **(C, D)** DIC and Fluorescence microscopy of mixtures of NBD-labeled NLs (DOPE_50_:DOPC_49.5_:NBD-DOPE_0.5_) and Rhodamine-labeled anionic liposomes ALs (DOPC_89.5_:DOPS_10_:Rho-DOPE_0.5_) prepared in water at R_AL/NL_ = 1 and incubated either with 2 % (w/v) 20 kDa PEO (D) or without PEO (C) for 6 hours at 25°C then imaged under DIC (left panels) and fluorescence microscopy (2nd, 3rd, 4th panels from left: NBD, Rho, and NBD/Rho combined channels).

For all remaining experiments (Figures 1B-5) we used NLs containing 50 mol% DOPE. We mixed NLs and fluorophore deficient anionic liposomes (ALs), consisting of 10 mol% negatively charged DOPS and 90 mol % zwitterionic DOPC, at a range of lipid molar ratios, R_AL/NL_ ((mol_DOPC (AL)_ + mol_DOPS (AL)_)/(mol_DOPC (NL)_ + mol_DOPE (NL)_)) between 1 and 8, in water and measured NBD fluorescence over time, then added 2% w/v 20 kDa polyethylene oxide (PEO) to the mixture after 26 minutes incubation. PEO is a known depletant which, when added into solution with larger objects, induces depletion attraction mediated clustering and phase separation of the objects and results in osmotic pressure on those phase separated objects proportionate to its concentration [56–59]. Consistent with other studies which used PEO to create liposome-EV hybrids [45], PEO driven depletion attraction and consequent membrane fusion drove NBD fluorescence to rapidly increase (Figure 1B, arrow). NBD fluorescence of the NL and anionic liposome mixture relative to unmixed NL increased proportionate to the increase in R_AL/NL_ following addition of 2 % w/v PEO (Figure 1B, right of arrow). This corresponds with visible clustering of NBD-labeled NL and Rhodamine-labeled anionic liposomes observed in differential-interference-contrast (DIC) and fluorescence microscopy immediately after addition of 2 % w/v PEO (Figure 1D). Prior to the addition of PEO, the NBD fluorescence of the NL and anionic liposome mixture relative to NL alone was constant (Figure 1B, left of arrow). This is likely because electrostatic repulsion of the net negative DOPS headgroup and the negative phosphate group on zwitterionic DOPE and DOPC in the opposing vesicle membranes in water (Debye length ∼1μm) was sufficient to prevent intermembrane distances from collapsing to the distance required for fusion to initiate. This is consistent with the observation that there was no visible clustering of NBD-labeled NL and Rhodamine-labeled anionic liposomes under DIC and fluorescent microscopy in water without depletant after up to 6 hours incubation (Figure 1C). Further, mixtures of NLs and anionic liposomes in high salt buffer (PBS, Debye length ∼1nm) show significantly more NBD dequenching relative to unmixed NLs compared to mixtures in water without PEO induced depletion attraction (Supplemental Figure 1). This supports the claim that electrostatic repulsion prevents fusion of NLs and anionic liposomes in this system given that the weaker electrostatic forces experienced by anionic liposomes in high salt buffer would lead to enhanced fusion because of charge screening. In contrast, the zwitterionic DOPC and zwitterionic DOPE and DOPC in opposing membranes of vesicles used in Figure 1A can arrange their dipole moments such that electrostatic repulsion is reduced so the intermembrane distance can collapse to the distance required for fusion, even in water and the absence of depletion attraction.

### PEO mediated clustering and osmotic pressure drives fusion of EVs and NLs

To compare fusion of NLs and EVs derived from distinct cell types (by FRET), we standardized the number of EVs in each mixture by measuring the concentration of EV particles through a Nanoparticle Tracking Analysis of each EV sample (Supplemental Figure 2). We maintained a ratio of 5nmol total NL lipid to 2×10^10^ EV particles in all experimental mixtures described here. The mixtures were made in PBS by combining the EVs and NLs, which was followed by obtaining an initial NBD signal measurement, then adding either water or 20kDa PEO in water to a final concentration of 2 % w/v PEO. Without the addition of PEO, the concentration of NL and EV particles was approximately 1.41 × 10^11^ particles/mL. At this particle concentration, the average ratio of NBD emission signal of the NL and EV mixture to unmixed NL was 1.31 ± 0.02 and 1.22 ± 0.04 for mixtures with PC3 and M21 cell derived EVs, respectively, after 150 minutes at 37°C (Figure 2 A and B, solid line curves). The addition of PEO results in depletion attraction mediated phase separation of the mixture of NLs and EVs which brings local particle concentrations of up to 1000—fold higher, at 1.52 × 10^14^ particles/mL assuming the EVs and NLs randomly packed and consisted of spheres with approximate diameters ≈200nm. The concentration of NLs and EVs at this density, along with osmotic pressure, resulted in an average NBD fluorescence ratio of 2.32 ± 0.18 and 2.23 ± 0.24 for mixtures of NLs and PC3 and M21 cell derived EVs, respectively, after 150 minutes at 37°C (Figure 2 A and B, short-dashed curves).

**Figure 2.**
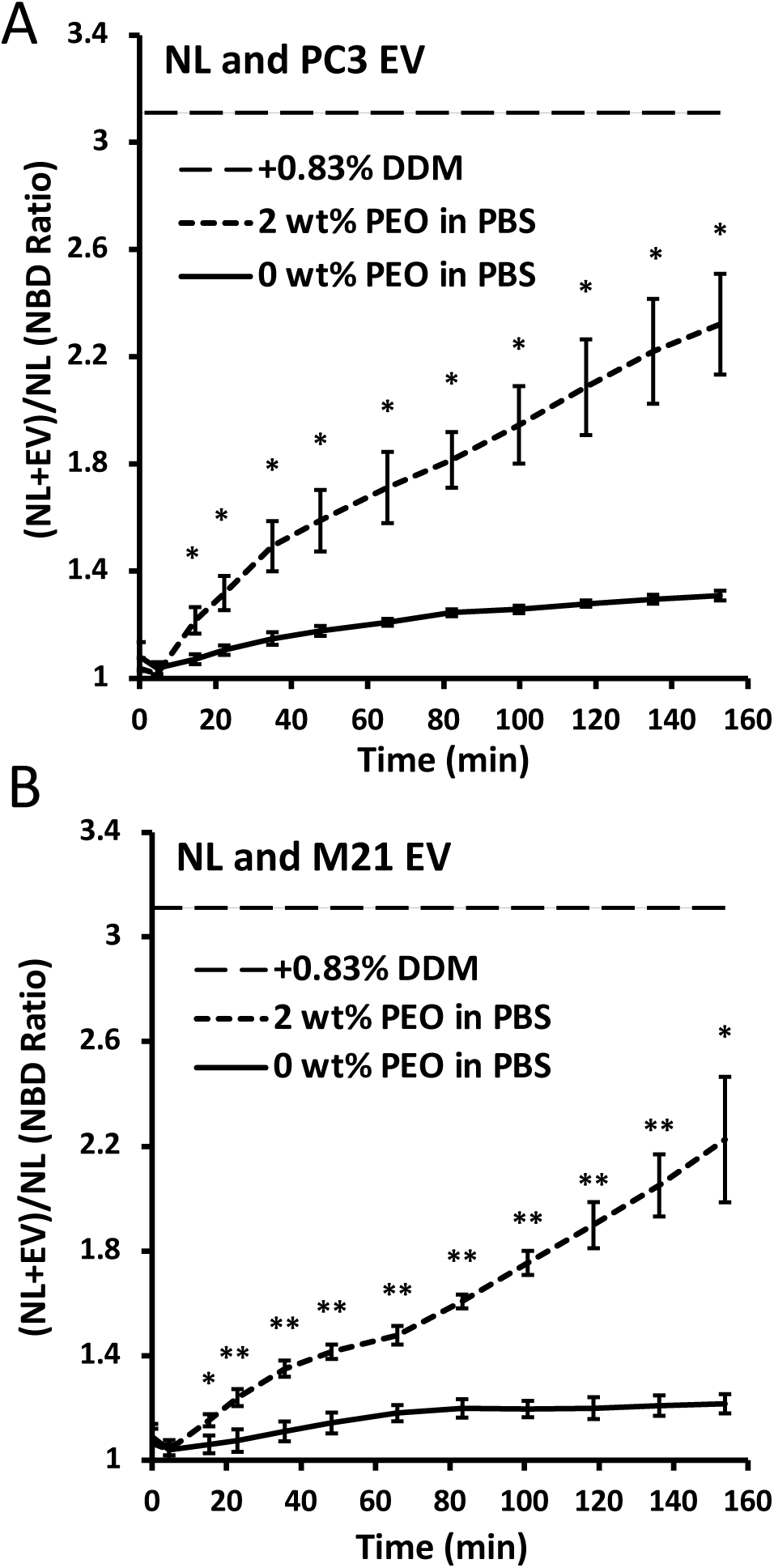
Total dequenched fluorescence (by FRET) of mixtures of neutral liposomes (NLs) and extracellular vesicles (EVs) with and without PEO. **(A, B)** PC3 (A) or M21 (B) derived EVs mixed with NBD and Rhodamine containing NLs (DOPE_50_:DOPC_49_:NBD-DOPE_0.5_:Rho-DOPE_0.5_) at a fixed ratio of 5 nmol NL lipid to 2 × 10^10^ EV particles in PBS, and at concentrations of 66.7 mM NL lipid and 1.33 × 10^11^ EV particles/mL, at 37°C with 2 % (w/v) 20 kDa PEO or without PEO. Total dequenching NBD fluorescence of these mixtures was taken over 150 minutes and normalized to the total NBD fluorescence of a sample of only NLs. The standard deviation of this NBD ratio was determined for three repeat samples and is shown as error bars at each data point. To quantify the maximum possible dequenching of the samples, dodecyl-beta-D-maltoside (DDM) was added to 0.83 % (w/w) in all mixtures of NLs and EVs and the average of these fluorescence values was normalized to samples containing only NL (long-dashed line at top). The standard deviation of the NBD ratio of these samples was 0.1 (not shown on chart).

To determine the theoretical maximum NBD ratio after full NBD dequenching, we used the detergent DDM to incorporate the liposomal lipids and fluorescent lipids into dispersed micelles and maximize the average distance between Rhodamine and NBD-labeled lipids according to established protocols [53]. The average NBD ratio of all mixtures upon DDM addition, compared to NL alone without DDM, was 3.11 ± 0.1 (Figure 2 A and B, long-dashed line). The quotient of the NBD ratio of NL and EV mixtures minus the starting NBD ratio of 1 divided by the average NBD ratio, obtained with DDM addition minus the starting NBD ratio of 1, gives the percent of maximum NBD dequenching achieved in each sample. Without PEO, EV and NL mixtures achieved 14.7% and 10.4% of maximum dequenching for PC3 and M21 cell derived EVs, respectively, after 150 minutes at 37°C. Mixtures of EVs and NLs which included PEO reached 62.6% and 58.3% of maximum dequenching for PC3 and M21 cell derived EVs, respectively, after 150 minutes at 37°C. These data demonstrate the efficacy of PEO mediated depletion attraction and vesicle phase separation in enhancing the fusion of EVs and synthetic liposomes.

### Acidic conditions enhance the fusion of EVs with NLs

Figure 3 (A and B) shows the result of our interrogation by FRET of the effects of acidity on fusion between PC3 and M21 cell derived EVs and synthetic NLs. We employed a discrete buffer system where mixtures were made in PBS or sodium acetate buffer and were pH adjusted to 7.4 and 5, respectively. We chose pH 5 as the acidic pH because the fusion and content release of EVs is documented at that pH, which is also just below the lower end of the pH range documented for late endosomes [4, 13]. The ratio of EV to NL was kept consistent with the PEO-mediated fusion experiments, at 5nmol total NL lipid to 2×10^13^ EV particles. NBD fluorescence measurements were taken immediately after the mixtures were made and taken at regular intervals over 150 minutes at 37°C. The average final NBD ratio of mixtures made in pH 5 buffer was 1.40 ± 0.04 and 1.44 ± 0.05 for PC3 and M21 cell derived EVs, respectively (Figure 3 A and B, dashed lines). The average final NBD ratio of mixtures made in pH 7.4 buffer was 1.31 ± 0.02 and 1.28 ± 0.01 for PC3 and M21 cell derived EVs, respectively (Figure 3 A and B, solid lines).

**Figure 3.**
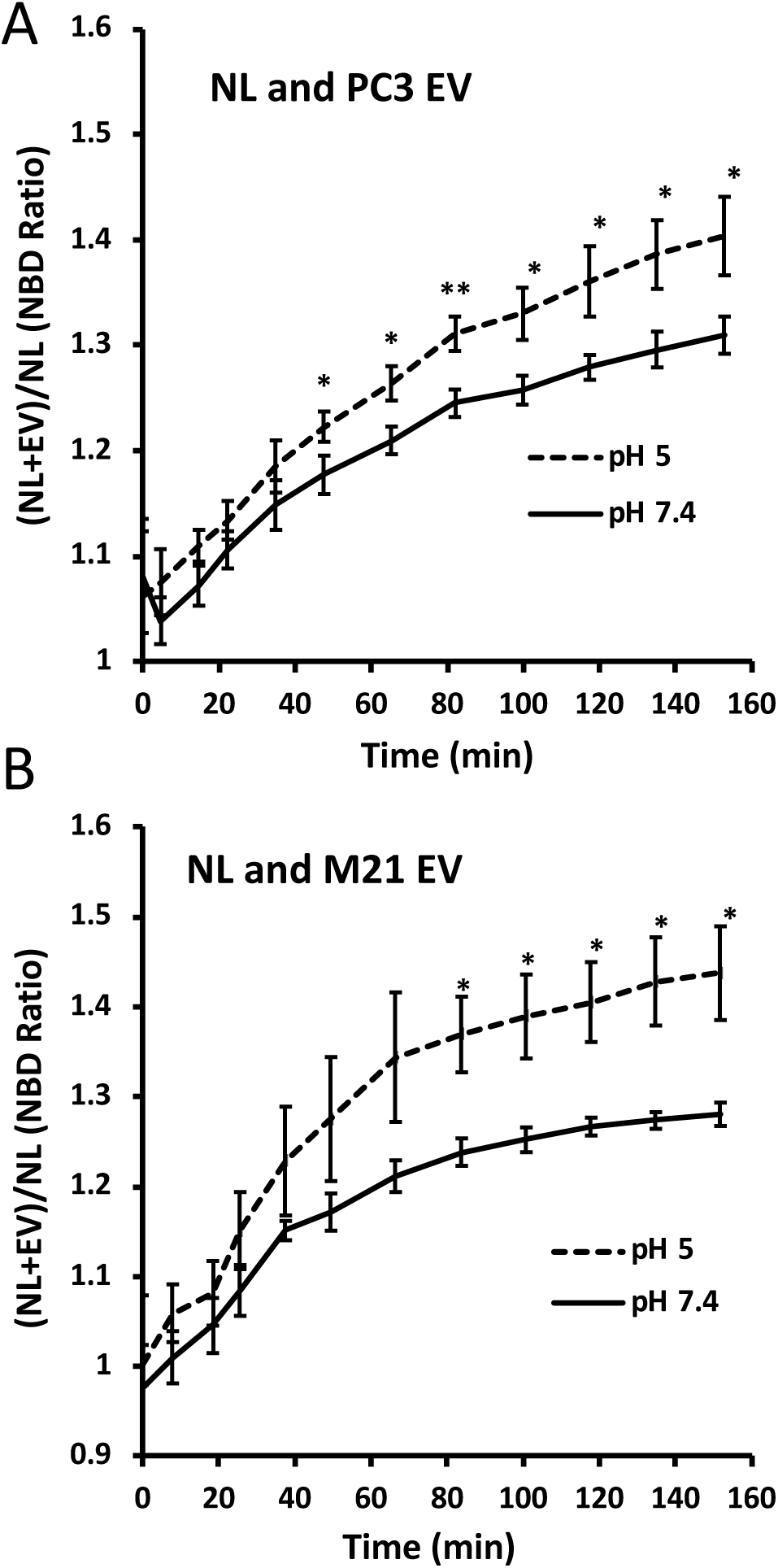
Total dequenched fluorescence (by FRET) of mixtures of neutral liposomes (NLs) and extracellular vesicles (EVs) in acidic or neutral conditions. **(A, B)** PC3 (A) or M21 (B) derived EVs were mixed with NBD and Rhodamine containing NLs (DOPE_50_:DOPC_49_:NBD-DOPE_0.5_:Rho-DOPE_0.5_) at a fixed ratio of 5 nmol NL lipid to 2 × 10^10^ EV particles in either sodium acetate buffer adjusted to pH 5 or PBS adjusted to pH 7.4 at 37°C. Total dequenching NBD fluorescence of these mixtures was measured over 150 minutes and normalized to the total NBD fluorescence of a sample of only NLs in conditions identical to the mixture being normalized. The maximum possible dequenching for these samples (not shown) was the same as that shown in Figure 2.

The difference between the final NBD ratio of mixtures made in pH 5 and pH 7.4 buffers was 0.09 and 0.16 for PC3 and M21 cell derived EVs, respectively. This was less than the difference between mixtures made with and without PEO, which was 1.01 for both PC3 and M21 EVs, indicating that the extent of fusion elicited by acidic conditions is less than that induced by depletion attraction mediated phase separation and osmotic pressure. Nevertheless, the enhancement of fusion, creating NL-EV hybrids under acidic conditions was reproducible and statistically significant across repeat experiments. Further, the observation that the absolute fluorescence of NBD in Triton X-100 treated mixtures prepared at pH 5 and pH 7.4 were 11341 ± 83 and 10788 ± 233, respectively, (data not shown) indicates that there were no pH-dependent effects on the fluorescence emission of NBD which could contribute to the difference in NBD ratio.

Taken together, the FRET data show that lipid-only synthetic liposomes show enhanced fusion with EVs, under physiologically relevant temperature and low pH conditions, with no other non-lipidic component, such as fusogenic amphipathic peptides, added to the synthetic liposomes. The FRET data also imply that NBD-lipid and rhodamine-lipid molecules have been delivered from the NLs to the NL-EV hybrid liposome in the fusion process.

### Acidic conditions facilitate clustering of EVs with other EVs and EVs with NLs under fluorescence microscopy

To obtain a mechanistic insight into the enhancement of fusion observed by FRET in mixtures of EVs and NLs, at low compared to neutral pH at 37°C, we performed direct imaging studies of the mixtures employing differential-interference-contrast (DIC) and fluorescence microscopy. Figure 4 shows DIC and fluorescent microscopy images of rhodamine-DOPE labeled EVs (Rho-EVs) and mixtures of Rho-EVs with NBD-DOPE labeled NLs in buffers adjusted to a range of pH values between 7.4 to 4.5. Without the addition of any synthetic liposome counterpart, we observed clustering of EVs under acidic conditions but not neutral conditions (Figure 4A, panels from left to right). Mixtures of rhodamine labeled EVs and NBD-DOPE containing NLs also clustered readily in acidic but not neutral conditions (Figure 4B, left versus right panels). The clustering of EVs in acidic conditions was similar to the clustering and phase separation induced by PEO-mediated depleted attraction (Figure 1D), suggesting that the enhancement of fusion may be in part due to the onset of short-range attractions between EVs or EVs with NLs at low pH. The formation of these clusters, specifically in acidic conditions but not neutral conditions, suggests that the surface properties of EVs are modified in acidic conditions such that the adherence of EVs to each other, and separately, to synthetic vesicles is promoted. This pH-dependent clustering of vesicles is observed in vesicles containing fragments of influenza virus hemagglutinin, which has a kink region that undergoes a structural shift at pH 5 to expose a hydrophobic domain to apposed membranes and promote clustering of adjacent vesicles [54]. Protein conformational changes are involved in the pH-dependent fusion and content release of EVs and cell membranes [13] and, given the precedence of this mechanism in other natural systems, may also be responsible for clustering of EVs and NLs and tethering of apposed membranes prior to fusion.

**Figure 4.**
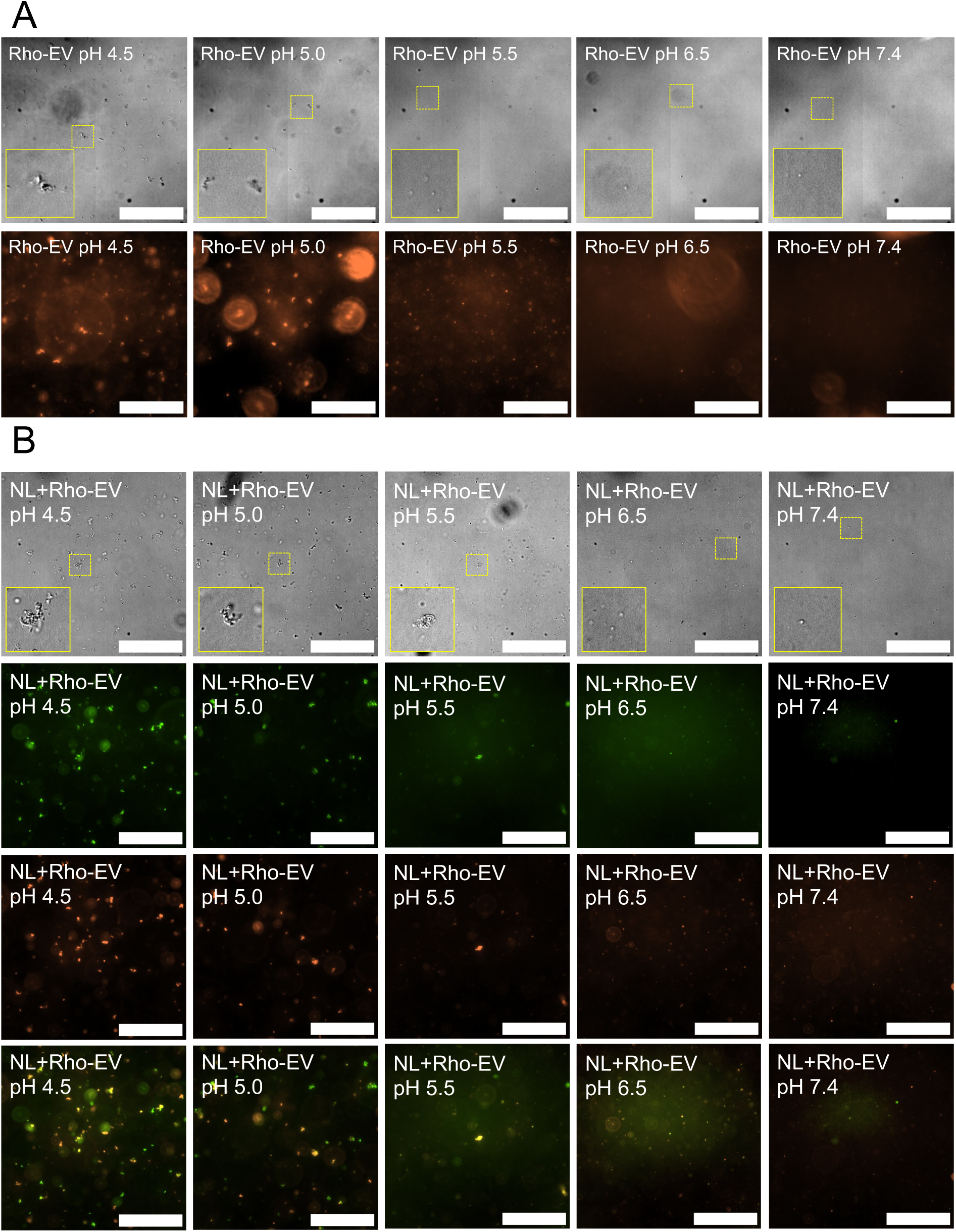
Differential-interference-contrast (DIC) and fluorescence microscopy of Rhodamine labeled extracellular vesicles (EVs) in the absence and presence of NBD labeled neutral liposomes (NLs). **(A, B)** DIC and fluorescence microscopy of Rhodamine-DOPE-labeled PC3 EVs alone (A) and in mixtures with NBD labeled NLs (DOPE_50_:DOPC_49.5_:NBD-DOPE_0.5_) (B) at decreasing pH values. EVs or mixtures of NLs and EVs were added to PBS pH adjusted to pH 7.4 and 6.5 by addition of HCl or sodium acetate buffer pH adjusted to pH 5.5, 5.0, and 4.5 by addition of HCl and incubated for 2 hours at 25°C then taken directly for DIC and fluorescence microscopy.

### Fusion of synthetic anionic liposomes and NLs are enhanced in acidic conditions

Enhanced fusion observed by FRET in EVs and mixtures of EVs and NLs in acidic conditions is likely the result of (i) surface protein modification, like the exposure of hydrophobic domains due to altered protein structure at low pH in enveloped viruses [55], (ii) altered electrostatics due to pH-dependent charge states in membrane phospholipid head groups, or a combination of the two. To investigate whether there is a contribution to enhanced fusion in acidic conditions from the charged membrane phospholipid head groups, we approximated the negatively charged phosphoserine content of EVs by using DOPS/DOPC containing synthetic liposomes as before and mixed these anionic liposomes (ALs) with NLs at R_AL/NL_=5 in pH 5 and pH 7.4 buffers. (Here, R_AL/NL_ = [(mol_DOPC (AL)_ + mol_DOPS (AL)_)/ (mol_DOPC (NL)_ + mol_DOPE (NL)_)].) We observed, in FRET, an NBD ratio of 1.77 ± 0.05 and 1.28 ± 0.04 for mixtures at pH 5 and 7.4, respectively, after 100 minutes at 37°C Figure 5A. The relatively high enhancement of NL and anionic liposome fusion at pH 5 compared to what is measured for NLs and EVs Figure 3 (A and B) suggests that there is a significant contribution to enhanced fusion of EVs and NLs at pH 5 from the phospholipid headgroups on EVs. Nevertheless, the lipid contribution to fusion (i.e. in going from the NL-AL to NL-EV mixtures) is attenuated by other membrane properties of the EVs. This may result from the steric stabilization of EVs by their surface proteins or the rigidity of their membranes decreasing the propensity for fusion at low pH despite crowding observed in those conditions.

**Figure 5.**
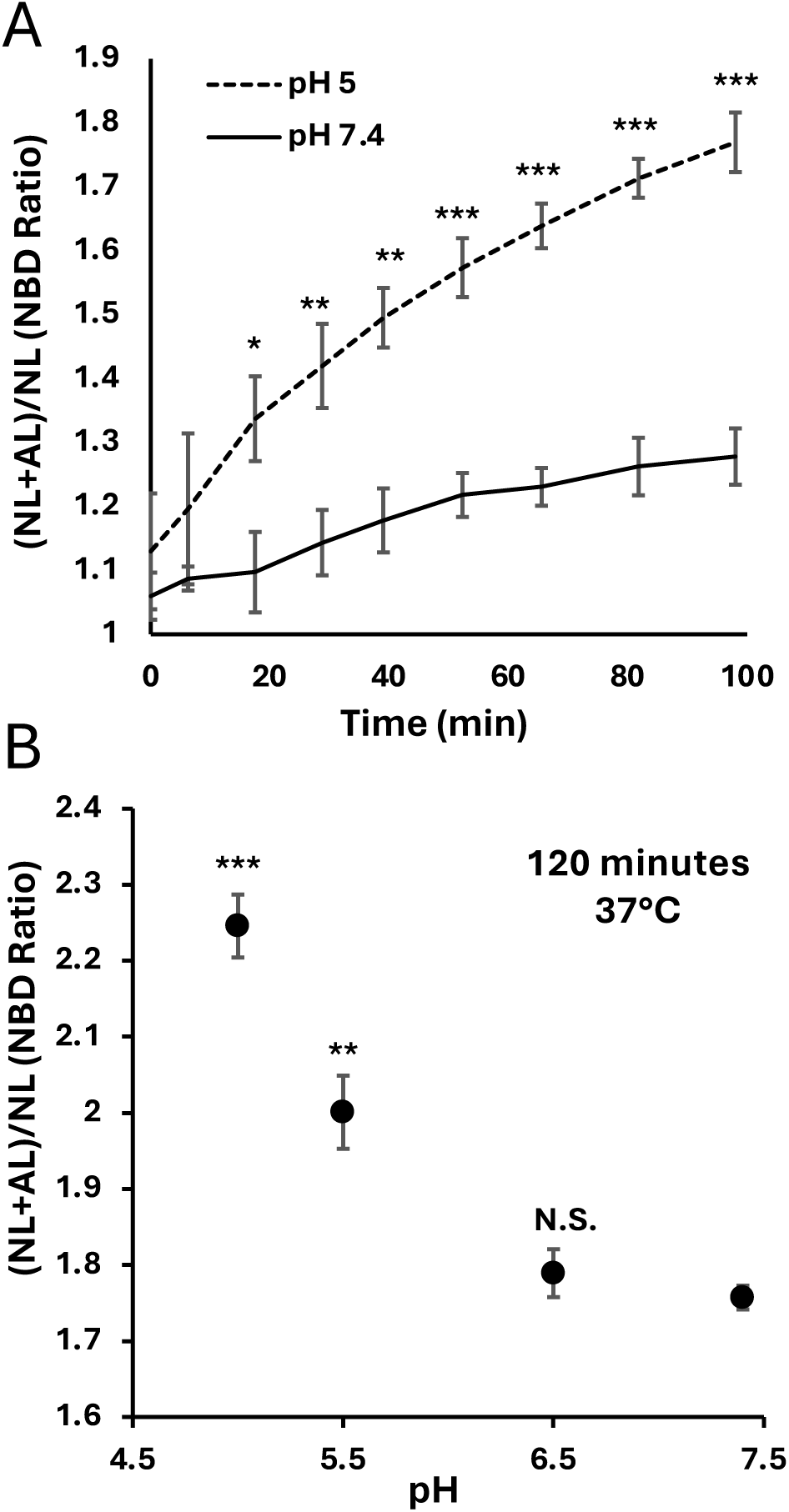
Total dequenched fluorescence (by FRET) of mixtures of neutral liposomes (NLs) and anionic liposomes. **(A, B)** Total fluorescence of mixtures of NBD and Rhodamine containing NLs (DOPE_50_:DOPC_49_:NBD-DOPE_0.5_:Rho-DOPE_0.5_) and anionic liposomes (ALs, DOPC_50_:DOPS_50_) over time at pH 5 and 7.4 (A) and after 2 hours across a range of pH values from 7.4 to 5.0 (B). NLs and ALs were mixed at R_AL/NL_ ((mol_DOPC (AL)_ + mol_DOPS (AL)_)/(mol_DOPC (NL)_ + mol_DOPE (NL)_)) = 5 in PBS or acetate buffer pH adjusted to 7.4 and 5, respectively, and held at 37°C while total dequenching fluorescence measurements were taken at discrete times over 100 minutes (A). NLs and ALs at R_AL/NL_ = 9 were mixed in PBS adjusted to pH 7.4 and 6.5 and sodium acetate adjusted to pH 5.5 and 5.0 then incubated for 2 hours at 37°C before total dequenched fluorescence measurements were taken. The results were plotted as the observed ratio of NBD dequenched fluorescence in mixtures of NLs and ALs at 2 hours (B).

It is important to note that the measured pKa value of phosphatidylserine, in the context of a liposome membrane also containing phosphatidylcholine in 100 mM salt concentrations, is 3.6 ± 0.1 [60]. Thus, our study reveals that significant impacts on the fusogenicity of anionic liposomes are observed well above the pKa of phosphatidylserine. To test whether the enhanced fusion at levels well-above the pKa of phosphoserine was a function of proton concentration, we varied the pH of mixtures of NL and AL at R_AL/NL_ = 9, in increments from 7.4 to 5 to determine the pH where enhanced fusion is first observed (Figure 5B). While little to no fusion was observed at pH 6.5, significant enhancement was observed at pH 5.5 and increased from there to pH 5, indicating that increases in proton concentration (by nearly two orders of magnitude between 7.4 and 5.5) drives the increase in fusion (Figure 5B).

Taken together, our observation of enhanced fusion between NLs and ALs at low pH 5.5 or 5 (but well above pH = pKa ≈3.6 where the carboxyl COO^−^ group on serine is protonated) suggests that more complex electrostatic effects, due to the two orders of magnitude increase in proton concentration between pH7.4 and 5.5, are responsible for the enhanced fusion effect rather than the relatively small protonation of phosphatidylserine headgroups (≈1.3 % to ≈3.8 %), which occurs at pH 5.5 and 5.

## CONCLUSION

Employing FRET in lipid mixing assays with established (NBD- and rhodamine-based) lipid FRET pairs that monitor the fusion of distinct vesicles, we show that acidic conditions enhance the fusion of EVs derived from prostate (PC3) and melanoma (M21) cancer cell lines with NLs compared to neutral conditions. The extent to which fusion was enhanced was less than what was observed with PEO but was repeatable and statistically significant. The fusion method is expected to spare the EV surface proteins from damage and leaves no residual PEO-depletant in the EV-liposome hybrid sample, enabling enhanced drug loading without loss of desirable targeting and biocompatibility properties conferred by EVs. The FRET data show directly that NBD-lipid and rhodamine-lipid molecules have been delivered from the NLs to the NL-EV hybrid vesicles created in the fusion process. We also show that acidic conditions drive clustering of EVs both with themselves and with NLs. This suggests a mechanism of EV fusion and content release which is like pH-mediated conformational change in influenza virus hemagglutinin where apposed membranes are anchored in proximity by polar tethers containing hydrophobic end-groups extending from the EV membrane similar to double-end anchored PEG lipids [61–63]. We further show that there is a component of enhanced EV fusion which is lipid driven, as fusion of synthetic anionic liposomes, which approximated the net negative charge and DOPS content of EVs, and NLs was enhanced to a greater degree than EVs and NLs at pH 5 compared to pH 7.4. The lower efficiency of fusion of EVs and NLs compared to anionic liposomes is likely due to the relative rigidity of EV, in particular, exosome membranes. Taken together, the findings of this study reveal that EV fusion is promoted in acidic conditions, and corresponds with concurrent clustering of EVs and adjacent vesicles, but requires additional interactions to achieve full fusion efficiency. This calls for the exploration of additional methods which leverage the negative charge of EV membranes or potential fusogenic EV membrane proteins in addition to the proximal anchoring achieved in acidic conditions. Building off this study, these weakly perturbative gentle methods would expand the therapeutic potential of exosomes by enabling high efficiency fusion of exosomes and liposomes containing therapeutic payloads to produce hybrid drug delivery vectors with targeting and biocompatibility superior to existing synthetic vectors.

## Supporting Information

FRET-based lipid mixing assay data of NLs and ALs used in Figure 1B incubated in either PBS or high resistivity water; nanoparticle tracking analysis results showing the concentration and size profiles of M21 and PC3 derived EVs as well as sonicated cationic liposomes.

## Notes

The authors declare no competing financial interest.

## ACKNOWLEDGEMENTS

The authors dedicate the paper to the memory of the biophysicist Erich Sackmann. The work was supported, in part, by the US National Science Foundation, Division of Materials Research, under award DMR-1807327. W.S.F. is grateful for a Connie Frank Fellowship Award to students conducting biomedical research for human health at UC Santa Barbara.

