## Supplemental Information for "Acidic Conditions Promote Clustering of Cancer Cell Derived Extracellular Vesicles and Enhance their Fusion with Synthetic Liposomes"

**Contents**

| FRET Data of NLs and ALs Incubated in PBS and High Resistivity Water | S1 |
| --- | --- |
| Nanoparticle Tracking Analysis of EVs and Cationic Liposomes. | S2 |

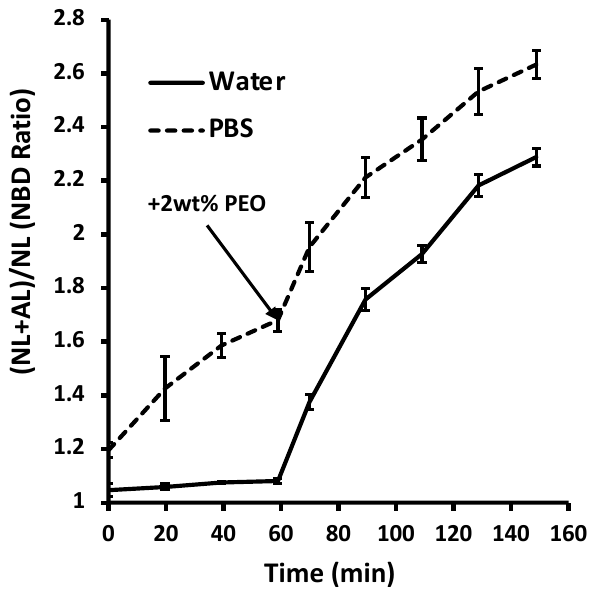
**FRET Data of NLs and ALs Incubated in PBS and High Resistivity Water**

**Supplemental Figure 1: Total dequenched fluorescence (by FRET) of mixtures of neutral liposomes (NLs) and anionic liposomes in water or PBS.** Total fluorescence measurements of NBD and Rhodamine containing NLs (DOPE_50_:DOPC_49_:NBD‑DOPE_0.5_:Rho‑DOPE_0.5_) and (ALs, DOPC_50_:DOPS_50_) mixed in water versus PBS. NLs and ALs were mixed at R_AL/NL_ ((mol_DOPC (AL)_ + mol_DOPS (AL)_)/(mol_DOPC (NL)_ + mol_DOPE (NL)_)) = 9 in water or PBS and held at 37°C without PEO then with PEO added to induce depletion attraction and osmotic pressure at 60 minutes. Total NBD dequenching fluorescence measurements of these samples were taken at regular intervals over 150 minutes. The fusion of NLs and anionic liposomes is enhanced by high salt conditions (PBS compared to water) in the absence of depletion attraction before the addition of PEO.

**S1**

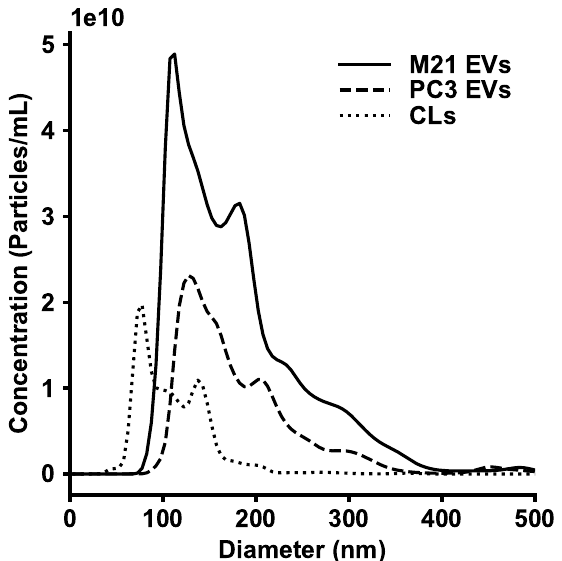
**Nanoparticle Tracking Analysis of EVs and Cationic Liposomes.**

**Supplemental Figure 2: Concentration and size profiles from nanoparticle tracking analysis (NTA) of extracellular vesicle (EV)** **isolates and sonicated cationic liposomes.** PC3 and M21 EVs isolated by serial centrifugation and ultracentrifugation as described previously were diluted 1:1000 into PBS and sonicated cationic liposomes (CLs) prepared as described in the methods section were diluted 1:1000 into water. These samples were taken directly for measurement using nanoparticle tracking analysis (NS300) and the measured concentration of particles with diameters ranging from 30 to 500 nm were plotted.

**S2**
